# Statistics in biology: a survey of the three major multidisciplinary journals

**DOI:** 10.1101/2025.02.04.636422

**Authors:** Jean Xavier Mazoit

## Abstract

This article presents the results of a survey of the three major multidisciplinary journals (Nature, Proceedings of the National Academy of Science, Science). Fifty articles involving experiments necessitating the agreement of an ethics committee were searched in each of the three journals to June 30th 2023. Because PNAs announced that a Statistical Review Committee was created and working since September 2023, a further set of 50 articles was retrieved and analyzed from 2024 January 1st. The following items were checked in articles: an explicit statement of approval from an animal ethics committee, the calculation of the appropriate samples size needed and how randomization and blinding were performed, the minimum sample size reported, the presence of repeats, and the methods used to limit type I and type II errors.

No clear experimental design was fully reported in any article. The major problems were 1) extremely small sample sizes (<4/group) in nearly half of the articles, 2) confusion between biological repeats and technical replications, and 3) lack of correction for multiple comparisons. These errors led to major inflation of type I and type II errors.

In conclusion, only 10 percent of the articles analyzed presented correct statistical methodology.

## Introduction

Statistics are notoriously wrong in biological journals including the major ones (1-3). While medical journals have regularly increased their standards in the past 50-60 years, biological journals have maintained the same low level of statistical analysis. The amount of money involved in health industry and in reimbursement of medical expenses as well as the pressure of regulatory agencies are likely a major cause of the rigorous approach in medical journals (4,5). Statistics courses are usually provided in medical schools and most medical journals have consultant statisticians who are systematically or on-demand involved in the reviewing process. Conversely, most biologists have a low background in statistics and even consider statistics negligible if not simply annoying. This attitude causes major mistakes in articles that may induce erroneous scientific and ethical conclusions. For example, a frequent mistake consists to sample one mouse, seed several plates and consider the data obtained as biological repeats rather than technical replicates (6). Experiments involving animals are particularly sensitive to these mistakes (7). This may pose ethical and scientific issues. Therefore, the present study was performed with the aim to quantify the most important statistical problems encountered in the three major multidisciplinary journals.

## Methods

Articles involving experiments on animals and necessitating the agreement of an ethics committee were searched in the three major multidisciplinary journals (Nature, PNAS [Proceedings of the National Academy of Science] and Science) to June 30th 2023. Although the aim of this study was only descriptive, the minimum number of articles to be analyzed in each group was calculated considering a two * three contingency table with an effect size (Cohen w) of 0.3 (medium), a power of 0.8 and an alpha risk of 0.0167 (0.05 corrected). Finally, because of possible exclusions, 50 articles/journal were analyzed. In addition, because PNAS established a Statistical Review Committee in September 2023 (8), an additional data set from January 1st 2024 was retrieved.

The following items were checked in articles:

1. The explicit statement of approval from an animal ethics committee, the reference to the ARRIVE checklist (9), the calculation of the appropriate number of animals or study units needed (power analysis) and how randomization and blinding were performed.
2. When between-group statistical comparisons were performed, the minimum number of subjects or units in each group, the presence of repeats (biological or technical), the methods used for analyzing results, and when multigroup analysis was performed, the methods used to limit type 1 error inflation were noted.
3. Finally, the report of statistics and if the actual p values were reported rather than a vague p<0.05.

In addition, because a great number of experiments were performed on very small samples, we calculated the power (1-β) of Student t tests (two-tailed) as a function of the number of subjects (equal number in each group) and of Cohen effect size d, using Gpower v. 3.1 for illustrative purpose (10).

## Results

### First data set (Fig. 1 and table 1)

Fifty consecutive articles involving animals were retrieved from 2023 March 15^th^ – June 28^th^ (Nature), from 2023 March 16^th^ – June 27^th^ (PNAS) and from 2022 November 25^th^ – 2023 June 30^th^ (Science). A total of 6 articles were excluded from analysis because their authors did not perform any statistical analysis or because the study was done on very small samples due to ethical reasons (Fig. 1). Less than 40 percent of articles presented almost correct statistical analyses (Nature n=12/47, PNAS n=18/49, Science n=18/48).

**Table 1.** Statistical analysis features in the three journals.

| <u>Ethics for animals</u> |  |  |  |  |  |
| --- | --- | --- | --- | --- | --- |
| Journal | Ethics committee * | ARRIVE | Power analysis ** | Randomisation | Blinding |
|  | Yes/No | Yes/No | Yes/No | Yes/Partly/No | Yes/Partly/No |
| Nature (n=47) | 47/0 | 1/46 | 2/45 | 18/8/21 | 11/10/26 |
| PNAS (n=50) | 46/4 | 2/48 | 0/50 | 2/0/48 | 1/1/48 |
| Science (n=49) | 49/0 | 4/44 | 2//47 | 13/6/30 | 8/10/31 |
| <u>Methodology for between group comparisons</u> |  |  |  |  |  |
|  | Minimum number of subjects/group | Test of normality | Adequacy of technical replicates | Proper analysis of longitudinal models |  |
|  | 2/3/4/>4 | Yes/No | in biological repeats |  |  |
|  |  |  | Yes/No | Yes/No |  |
| Nature (n=47) | 0/25/11/11 | 10/37 | 1/13 | 4/13 |  |
| PNAS (n=50) | 0/17/9/24 | 6/44 | 1/6 | 1/13 |  |
| Science (n=49) | 1/20/9/19 | 8/41 | 0/13 | 6/8 |  |
| <u>Correction for multiple comparisons</u> |  |  |  |  |  |
|  | FWER correction # | Adequate use of FDR † | Logrank test |  |  |
| | Yes/No | Yes/No | 2 groups | > 2 groups \$ | |
| Nature (n=47) | 23/18 | 26/1 | 10 | 7 |  |
| PNAS (n=50) | 27/17 | 2/6 | 2 | 4 |  |
| Science (n=49) | 22/20 | 9/3 | 4 | 3 |  |

\* Clear mention of approval by an ethics committee; \*\* Power analysis: calculation of the minimum sample size needed for statistical analysis;

**Figure 1.**
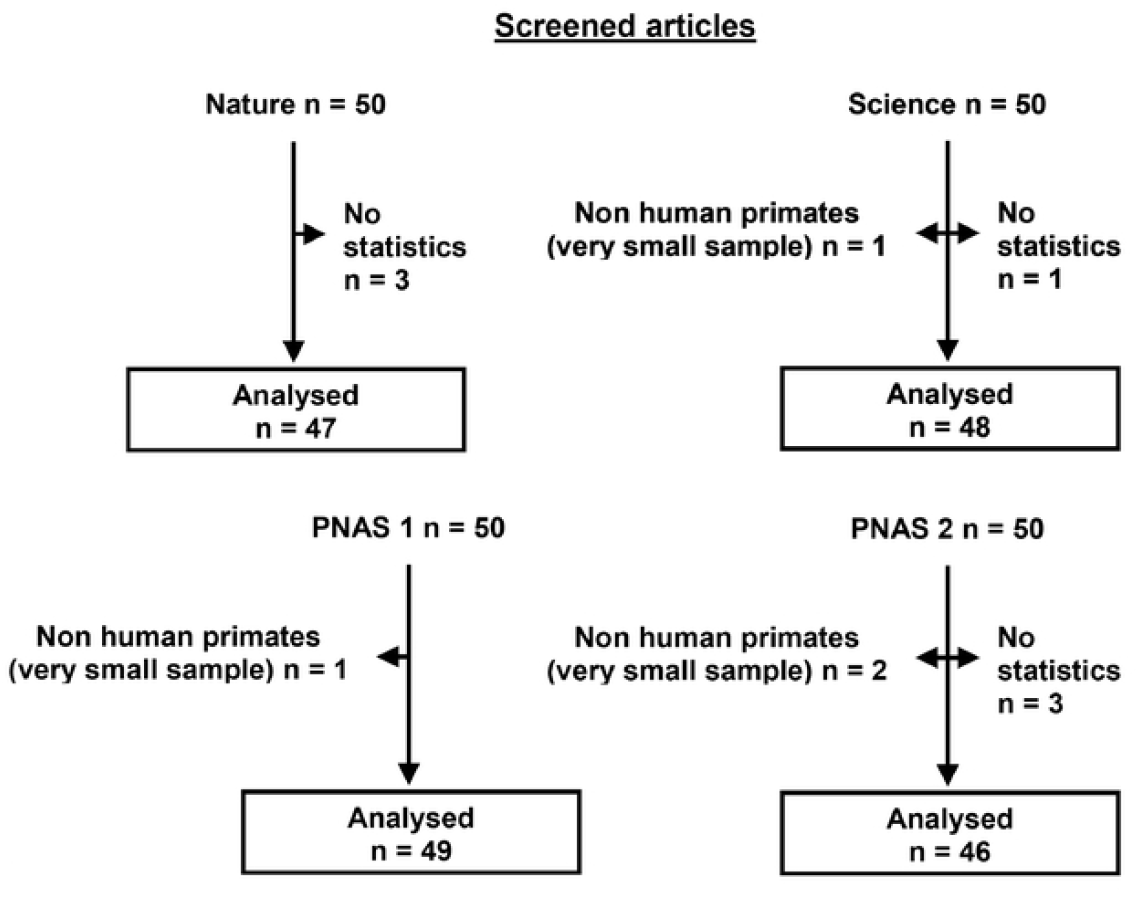
Flowchart of the study.

### Animal experiments and design

No clear experimental design was reported in any journal. Similarly, less than 10% ARRIVE statement was cited and sample size calculation was mentioned in less than 10 % of articles. Randomization and blinding were performed mostly for microscopic slides reading, and performed in one third of the articles.

### Between group comparisons

The two major problems were extremely small sample size and confusion between biological repeats and technical replications.

A test of normality was not performed in the vast majority of studies. The minimum number of subjects per group was less than four in 55 %, 37 % and 48 % of the articles in Nature, PNAS and Science respectively. In addition to impair the validity of the comparison (type I error), these very small sample sizes (2 subjects/group in three cases) led to a major risk of type II error. Power as a function of sample size and effect size is reported in Fig. 2, which shows that a power of 0.8 is far from being reached with samples sizes of 3 subjects/group. In addition, when one consider reasonable values of Cohen d, i.e. less than 4, only a minimum of 5 subjects/group allows to reach a power of 0.8; and even then, only with an alpha error set at 0.05. As a consequence, experiments with less than 5 subjects/group have a very high rate of false negative.

**Figure 2.**
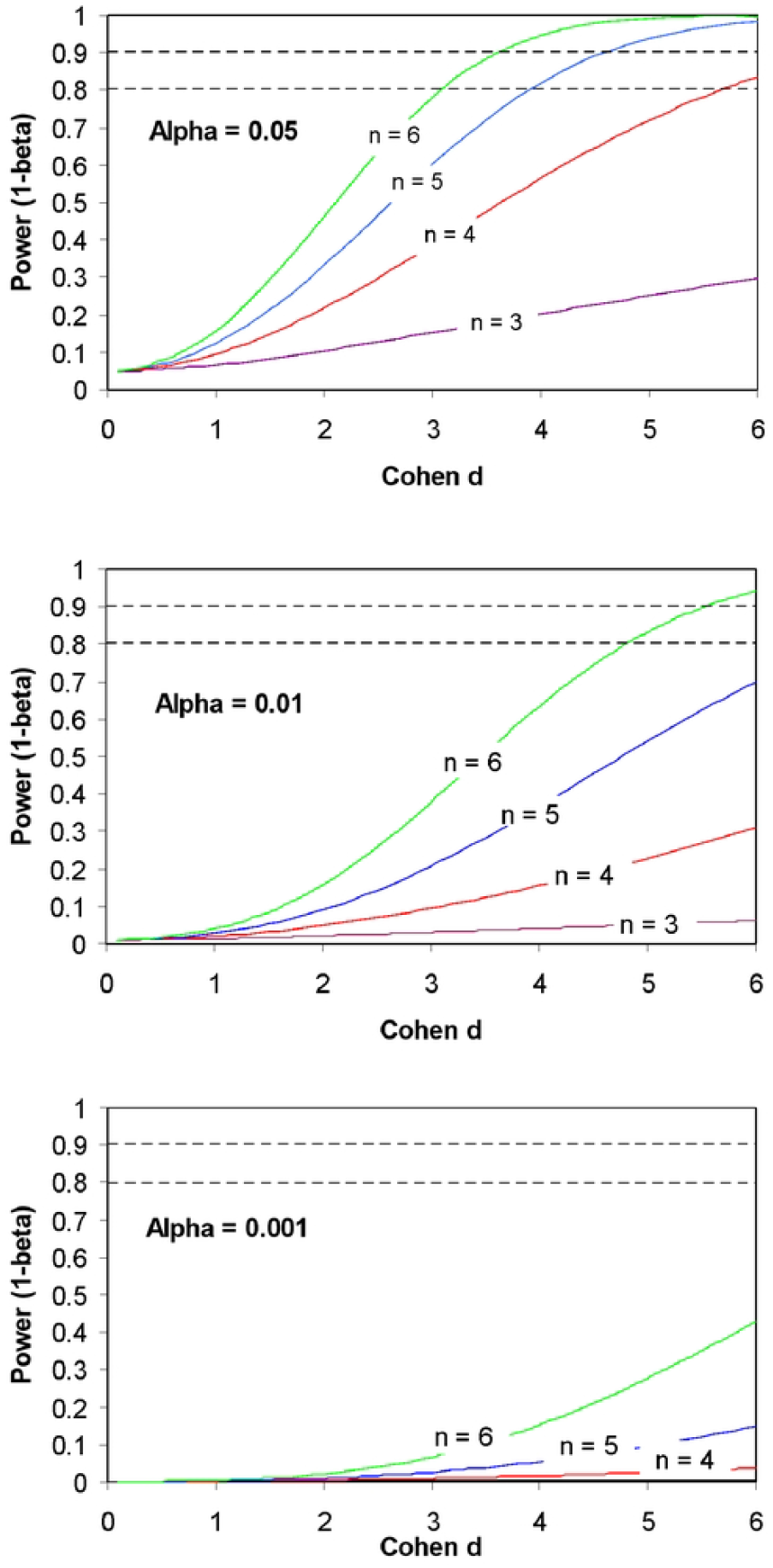
Power of a Student t test (two-tailed) as a function of effect size (Cohen d) and sample size (n = 3 to 6). Three conditions of type I errors (alpha) are represented: 0.05, 0.01, 0.001. The two dashed lines represent 80% and 90% power. Clearly, with reasonable values of effect size (Cohen d ≤ 4), less than 5 subjects/group is insufficient to attain a power of 80%. Even with these conditions, the problem of type I error remains. Data were generated using Gpower 3.1.

### Repeats

Repeats and replicates are most often misunderstood. This confusion between technical and biological repeats was noticed 14 times in Nature, 15 times in Science and 12 times in PNAS. In addition, proper analysis technique (i.e. nested ANOVA or mixed-effect model) was noted only 13 times.

The absence of correction for multiple comparisons was noticed frequently. In addition, False Discovery Rate (FDR) correction was sometimes used in place of Family-Wise Error Rate (FWER) correction to compare very small samples. Survival analysis was followed by a logrank test in nearly all cases. The test always lacked continuity correction in case of extremely small samples and correction for multiple comparisons when more than two groups were compared.

Repeated measures are another issue. For example, when the same mouse is observed several times, the correlation between these measures was not taken into account. This problem was observed in 13, 5 and 12 articles in Nature, PNAS and Science respectively.

### Additional data set (Fig. 1 and table 1)

Fifty consecutive articles published in PNAS involving experiments in animals were retrieved from 2024 January 1st – 31st. A total of five articles were excluded from analysis because their authors did not perform any statistics (three articles), or because the studies involved non human primates with a limited sampling size (n=2) due to ethical reasons (two articles) (Fig. 1). Only twenty of the remaining 45 articles presented almost correct statistical analyses.

Unfortunately, despite the establishment of the committee, the problems were similar (item PNAS2, table 1).

## Discussion

The present survey shows that improper statistical analysis is common in life sciences. This diagnostic is not new (1-3, 8); however the extent of the problem appears of utmost importance. Indeed, this poses ethical issues, not only because of the problem of inadequate animal testing, but also because of the great impact these articles published in major journals may have on societal choices and future orientations of applied research.

Animal experiments lack previous calculation of the right sample size and complete randomization and blinding are never included in a proper experimental design. The paradox is that the choice of too small samples is often favoured by ethics committees. Researchers multiply the experiments at the expense of the quality of research. An example is the multiplication of technical replications, which are often believed to be biological repeats (this mistake was noticed in 75 % of the 63 cases of explicit replicates reported). Together with analysis on extremely small samples (less than 5 subjects/group), this leads to major type I and type II error inflation.

### Type I errors

Samples of extremely small size (less than 5 subjects/group, see table 1) whether analyzed using parametric or non parametric tests is a key issue (11,12). Some authors are aware of the problem and perform repeats. However, the confusion between biological repeats and technical replications makes the problem worse (Table 1) (6). By performing repeats, most authors believe that this ascertains the robustness or their results. This is wrong. Firstly, they usually only reproduce their experimental bias. A proper repeated experiment should be performed in an independent laboratory, far from the initial one. Secondly, it is important to note that the probability that a second experiment reach also statistical significance is lower than expected by the researchers (13,14). For example, the probability that a second experiment reach p=0.05 after an initial experiment significant at p=0.05 is only 0.5; and after a first p=0.01 experiment is approximately seven in ten. In addition, what if the second experiment is not significant ? To pool the data ? Certainly not ! In this case, it is better to perform an analysis with an inter-occasion variability parameter. However, an initial power analysis leading to a moderately greater sample size should have been a much better choice. This leads to more robust results with less subjects (animals).

The rational of technical replications, for example in culture plates, is to synthesize the data in a statistic like the mean or the median, and then to decrease the measurement error. However, often there is no clear boundary between biological and technical repeats. Thus, we made the following choice: cell lines or cells dissociated for the purpose of the study (dissociated neurons from embryos for example) have be considered as a population in which samples were taken. Conversely, repeated seeds from one mouse are clearly technical replications and in any case biological repeats.

In this regard, there is a semantic confusion between experimental and statistical samples, even in forms that the authors must fill when submitting their manuscript. This brings us to the problem of study design. A clear experimental plan should be designed before the start of experiments and described in the report.

### Type II errors

When comparing very small samples, power of the test is extremely low (Fig. Together with under publication of negative results, this inflation of false negative rate is a major bias in research in general. In addition, the “winner’s curse” effect inflate the magnitude of the effect size when a “significant” effect is discovered (3,15). Fortunately, when the experiment is repeated by another team, the effect size decreases because of regression to the mean (16). This is another reason to recommend repeats, but in other, independent laboratories.

Corrections for multiple comparisons are often ignored or wrongly considered. Because researchers want to describe each tree, rather than the whole forest, all possible post-hoc tests are performed after an analysis of variance (ANOVA). In this case, a non corrected Student t test is the choice in almost half of the articles (table 1). Some authors used the False Discovery Rate (FDR) correction in place of Family-Wise Error Rate (FWER) correction to compare very small samples. Procedures to control FDR have been conceived for large-scale testing and they are exploratory rather than confirmatory (17). Log rank tests also suffer from the same lack of correction for multiple comparison as well as for very small samples comparisons (18). In one case, the authors deliberately did not correct the risk, but let the reader judge by giving the raw probabilities. In the same way, it is recommended to give the full probability rather than a vague p less than 0.05 (19). Recently, some researchers have recommended considering statistical significance at 0.005 (20). However, in the eternal debate between the Ronald Fisher and Jersy Neyman approach, we are obliged to be pragmatic: it appears mandatory 1) to perform a power analysis before initiating the study, but report full probability values and 2) to increase the size of samples - less than five subjects/group should be an exception.

At last, longitudinal models, i.e., models in which the same subject is observed more than once are inappropriately analyzed three times out of four. In these models, repeated observations are correlated and the number of degree of freedom is, therefore, less than in a classical ANOVA.

The use of mixed-effect models may solve many problems. These models correctly and easily apprehend technical replicates nested in biological repeats, repeated measures and importantly, unbalanced data sets.

A second data set was retrieved four months after the announcement in PNAS that a Statistical Review Committee was created and working. Unfortunately, the situation is similar as before, likely because reviewers - who are supposed to seize the committee, and authors have the same lack of statistical knowledge.

In conclusion, only 10 percent of articles analyzed present appropriate statistical analysis. To remedy this problem, it may be recommended that consultants in methodology or statistics review most, if not all, articles before publication. Indeed, the problem will not be solved by the wave of a magic wand. This may be a difficult and costly task, but medical journals have been able to meet the challenge. It is a question of credibility for the “great” journals.

